# Protein Ensemble Generation through Variational Autoencoder Latent Space Sampling

**DOI:** 10.1101/2023.08.01.551540

**Authors:** Sanaa Mansoor, Minkyung Baek, Hahnbeom Park, Gyu Rie Lee, David Baker

## Abstract

Mapping the ensemble of protein conformations that contribute to function and can be targeted by small molecule drugs remains an outstanding challenge. Here we explore the use of soft-introspective variational autoencoders for reducing the challenge of dimensionality in the protein structure ensemble generation problem. We convert high-dimensional protein structural data into a continuous, low-dimensional representation, carry out search in this space guided by a structure quality metric, then use RoseTTAFold to generate 3D structures. We use this approach to generate ensembles for the cancer relevant protein K-Ras, training the VAE on a subset of the available K-Ras crystal structures and MD simulation snapshots, and assessing the extent of sampling close to crystal structures withheld from training. We find that our latent space sampling procedure rapidly generates ensembles with high structural quality and is able to sample within 1 angstrom of held out crystal structures, with a consistency higher than MD simulation or AlphaFold2 prediction. The sampled structures sufficiently recapitulate the cryptic pockets in the held-out K-Ras structures to allow for small molecule docking.

## Main Text

A major challenge in drug discovery is identifying cryptic binding pockets that can be targeted by small molecule drugs (Beglov et al., 2018; Vajda et al., 2018; Vijayan et al., 2015). Despite considerable advances in single state native protein structure prediction with AlphaFold and RoseTTAFold in the past several years, generating plausible ensembles of structures that can be populated upon binding a small molecule, or during protein function, remains an outstanding problem– AlphaFold and RoseTTAFold generate single structures, rather than ensembles. Molecular dynamics (MD) trajectories generate protein ensembles by simulating protein motion around the native structure, and are often used to generate ensembles prior to small molecule docking calculations, but often fail to identify cryptic ligand binding pockets not present in the unbound structure (Cimermancic et al., 2016; Beglov et al., 2018; Vajda et al., 2018; Vijayan et al., 2015) or require very long and hence highly compute-intensive simulations (typically sub-to-several microsecond level) (Kimura et al., 2017; Meller et al., 2023; Sun et al., 2020). Other classical approaches have been used to sample protein conformers through Rosetta (Larson et al., 2002), and loop sampling using kinematic closure (KIC) (Mandell et al., 2009), but have not sampled the types of conformational changes involved in cryptic pocket formation. On the deep learning side, variational autoencoders which project complex data into a smaller dimension latent space have been used to generate alternative backbones for general protein design tasks such as de novo design of 64 residue backbones (Anand et al., 2018), graph-based protein design (Ingraham et al., 2019) and Ig-fold modeling (Eguchi et al. 2020). VAEs have been used previously to sample the conformational space of proteins, but have required visual inspection of the trained latent space to sample (Tian et al., 2021), or have focused on mapping correlative fluctuations in extensive MD simulations of both the apo and holo state of a target protein (Tsuchiya et al., 2019).

We reasoned that sampling within the latent space of variational autoencoders could provide a solution to the ensemble generation problem for a specific protein sequence. Unlike most previous VAE approaches, which have trained on many different proteins, the challenge of a protein specific VAE is that there is limited training data. We reasoned this limitation could be overcome by supplementing available crystal structures of the protein of interest in alternative conformations with snapshots from short MD trajectories started from each of these structures. For exploring this approach, we chose the critical cancer target K-Ras as a model system due to its considerable therapeutic importance and the many available structures (Pantsar et al. 2019; Liete et al. 2022).

We began by exploring different VAE architectures, training on ensembles of MD simulations from alternate crystals forms of K-Ras (full details in the Methods section), and evaluating the quality of 3D reconstruction following encoding and decoding. For encoding 3D structural information, we chose to use the two-dimensional RoseTTAFold (Baek et al., 2021) template features. The reconstructed template features were then used as input template features for 3D structure generation with RoseTTAFold, along with the amino acid sequence. We evaluate the accuracy of reconstruction by computing the RMSD between the input and output 3D coordinates. The RMSD loss was calculated based on the 2D template features of the input and output 3D structures. We generate new samples by guided exploration in the latent space, followed by 3D coordinate generation with RF (RoseTTAFold, Figure 1).

**Figure 1.**
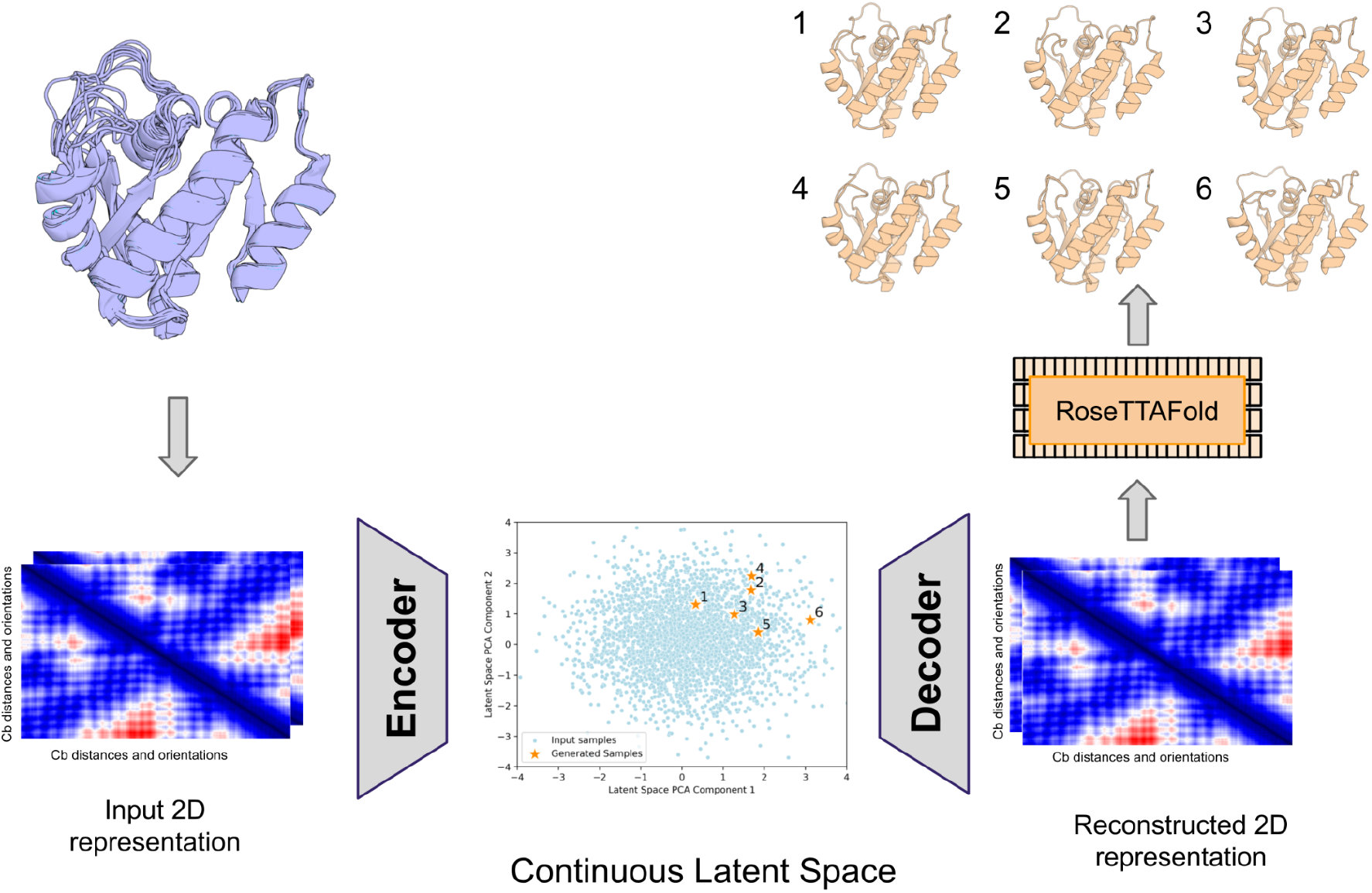
VAE based ensemble generation approach. 3D coordinates from crystal structures and MD simulations are converted to RoseTTAFold 2D template features (Baek et al., 2021). The decoded template features are converted to 3D structures through RoseTTAFold, which is also given the amino acid sequence. Ensembles are generated by sampling in the latent space followed by decoding and RF structure generation.

The reconstruction accuracy of crystal structures not included in the training set provides a rough lower bound on the accuracy with which our approach can recapitulate conformations of interest. For each available K-Ras structure, we trained a VAE leaving out this structure and others within 1A RMSD, and evaluated the accuracy of reconstruction following RoseTTAFold (Baek et al., 2021) 3D coordinate generation. We obtained best results with the soft-introspective VAE architecture (Figure S1), and the accuracy of reconstruction plateaued at ∼256 latent space dimensions (Figure S2) For most of the targets (13/20), the reconstructed target from the VAE was of sub-angstrom accuracy (RMSD < 1A); for comparison only 2/20 AF2 structure predictions were of sub-angstrom accuracy (Figure 2).

**Figure 2.**
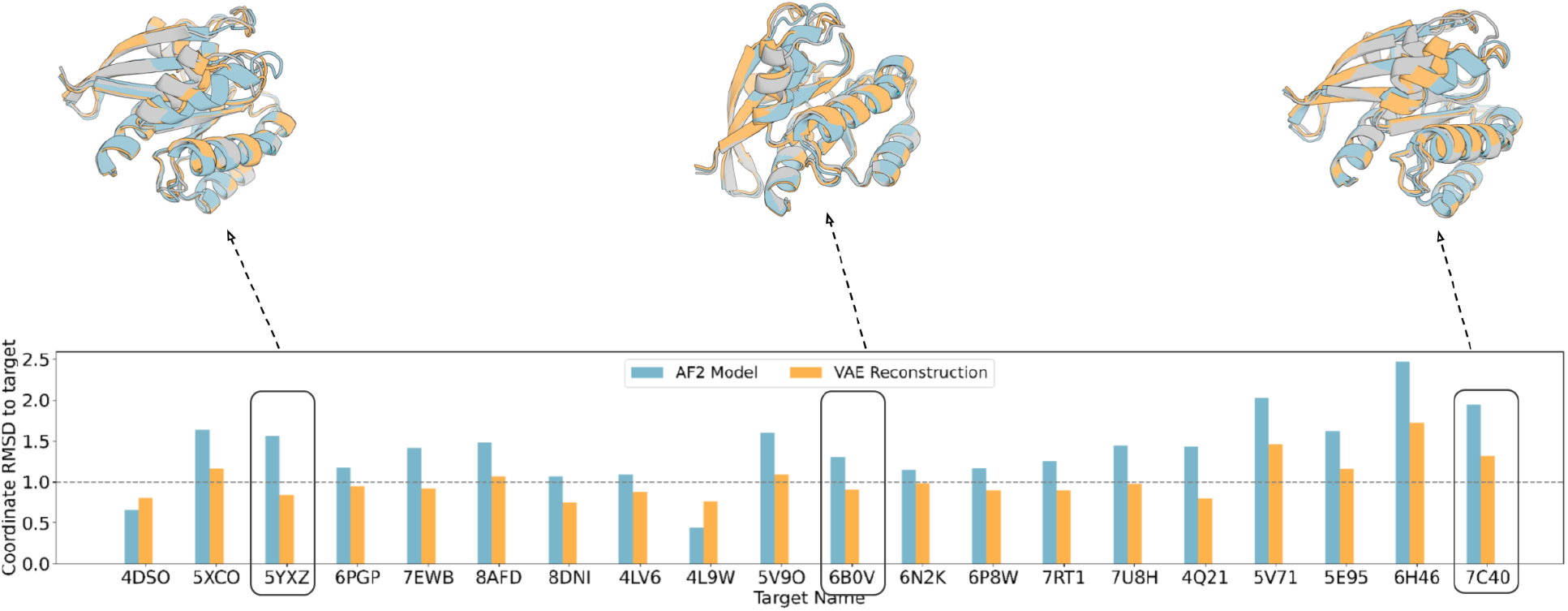
VAE Structure reconstruction accuracy. Coordinate RMSD of the closest AF2 predicted model and the reconstructed model from the VAE decoded template features generated using RoseTTAFold. Structural superimpositions for 3 targets are highlighted on top with the target crystal in gray, the AF2 prediction in blue and the VAE reconstruction in orange.

We next explored the possibility of generating plausible K-Ras ensembles by sampling in the latent space of the trained VAEs. To help ensure that the sampled structures remained broadly consistent with the sequence and were physically plausible, we guided sampling by the consistency to the AF2 predicted distance distribution for the amino acid sequence. Samples were generated from a normal distribution with a mean of 0 and variance of 1, decoded into the corresponding Cb distance map, the CCE to the AF2 predicted distogram for the sequence was computed, and local optimization in the latent space was carried out through gradient descent on the CCE value, limiting the total (latent space) distance traversed from the starting point to prevent convergence. Following decoding and RF structure generation, samples were evaluated using coordinate RMSD to the target crystal on either the overall structure recapitulation and cryptic pocket environment reconstruction (defined as the residues within 5 angstroms of the ligand binding pocket).

Using this VAE guided sampling approach, we generated K-Ras structure ensembles, again holding out individual K-Ras crystal structures and MD simulation snapshots derived from them, along with other K-Ras crystal structures (and MD snapshots) within 1 angstrom RMSD. We evaluated these ensembles by determining how closely they sampled the held out structures. An advantage of our approach is that ensembles can be generated quite rapidly (compared to MD simulations, for example), and the closest RMSD to the held out structures of course decreases with increasing number of samples (Figure S3). We found that ensembles of 3000 structures sampled more closely to the held out structures than the closest training set crystal structure, training set MD simulation snapshot and the closest AF2 model for most targets (Figure 3).

**Figure 3.**
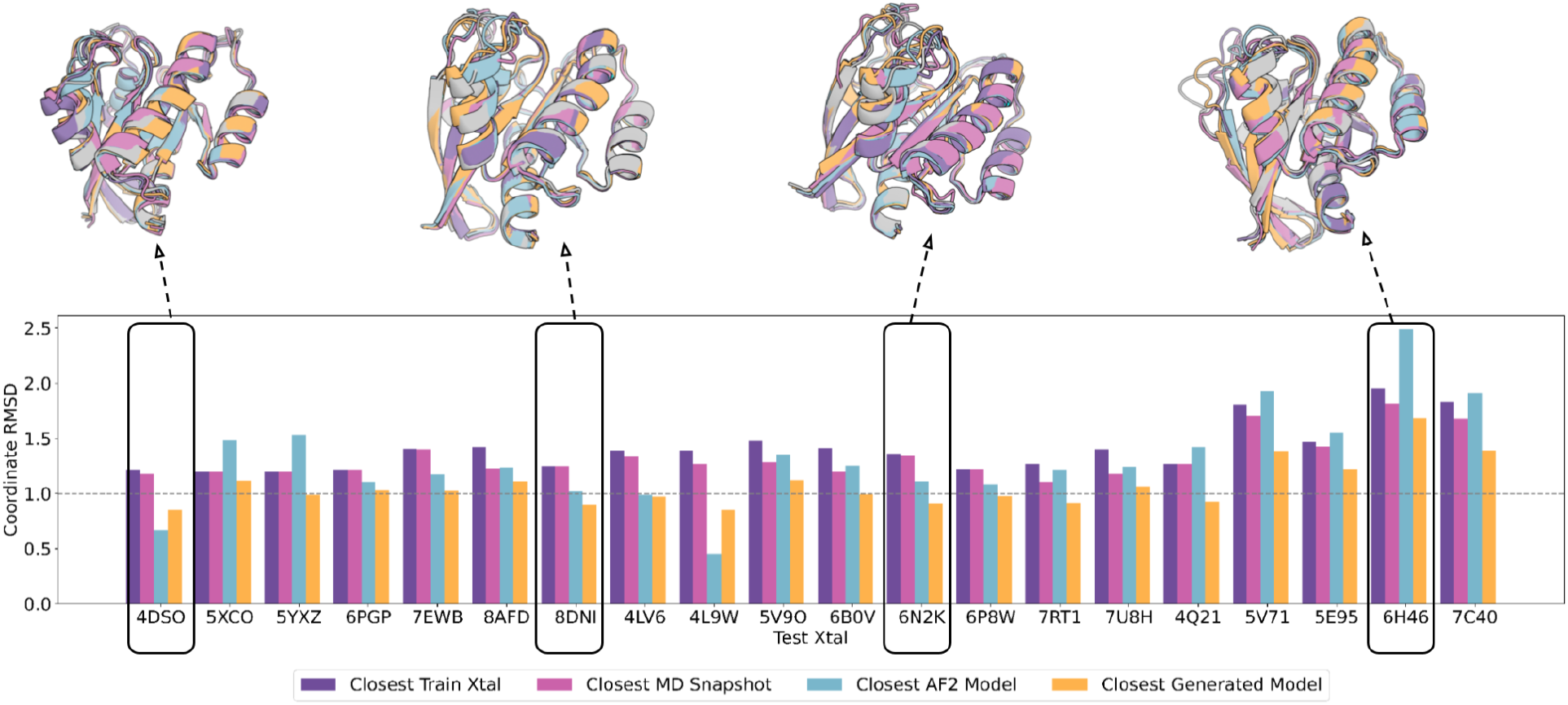
The VAE enables sampling closer to held out K-Ras crystal structures than MD or AlphaFold generated structures. For each test crystal structure (name below bars), a VAE was trained using MD simulation data from all crystal structures with greater than 1A RMSD, and used to generate a structure ensemble. Bars indicate the coordinate error to the test crystal of the closest train crystal, the closest training sample, the closest AF2 model and the closest VAE generated sample.

For small molecule docking calculations, the sampling of alternative ligand binding pocket geometries is particularly important. Comparison of the RMSD over the ligand binding pocket residues between the closest sampled conformation in the generated ensembles and the held out structures showed that the ensembles sample closer than the closest training MD snapshot or crystal structure in most cases (Figure 4). Structural superimpositions show that the generated samples do not clash with the superimposed ligand from the target structure, highlighted in orange, and therefore can be docked without hindrance, whereas for the closest train crystal and the closest AF2 model, there are significant clashes (Figure 4).

**Figure 4.**
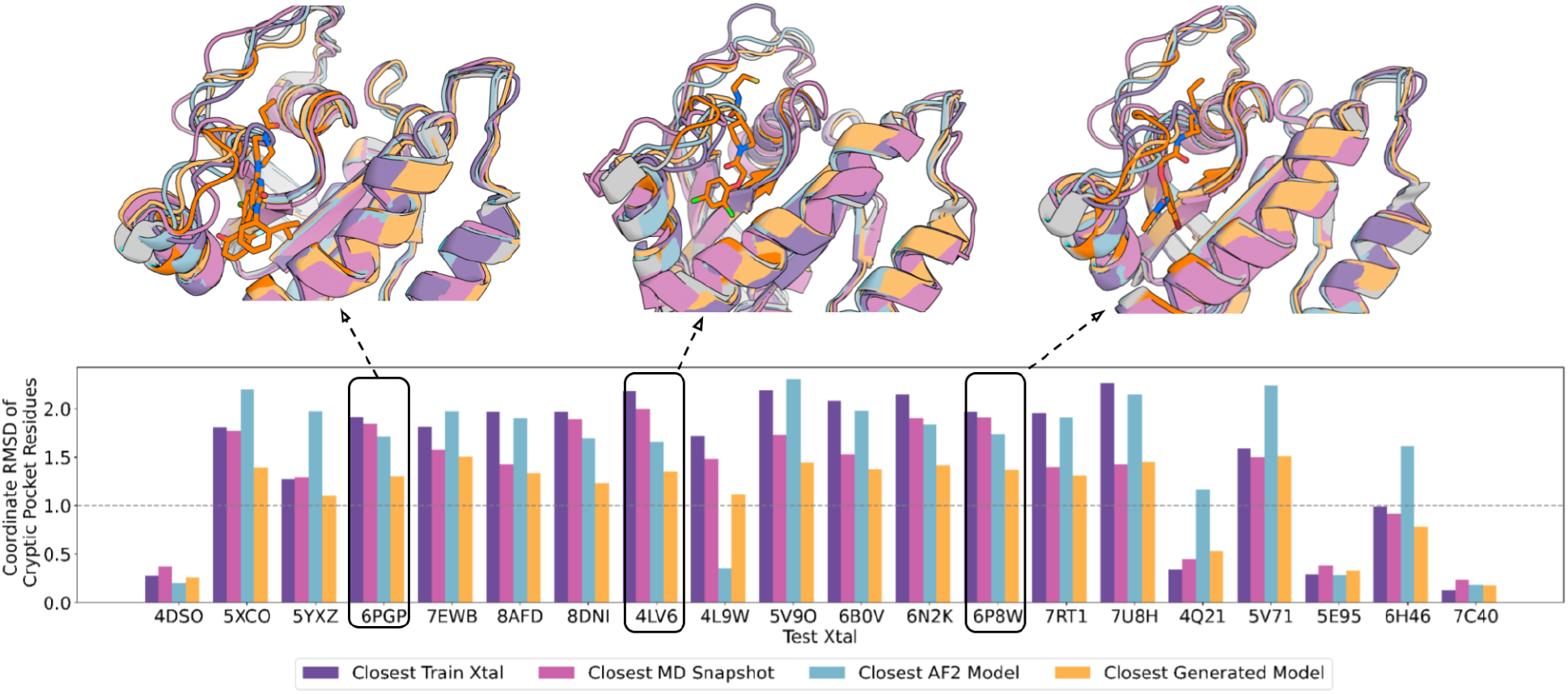
VAE sampling of K-Ras cryptic pocket geometry. As in Fig 3, but with the coordinate error to the test crystal structure computed over only the binding site residues (defined as the residues within 5 angstroms of the ligand binding pocket). The structural superimpositions (top) show the ligand inhibitor docked in only the target crystal, where the cryptic binding pocket and the ligand are highlighted in orange on the target crystal structure.

We used the physically based GA-ligand docking method to dock ligands onto all the models generated from the VAE, the training examples and the AF2 models. Consistent with the above observations, the RMSD over the ligand atoms was consistently lower for the ensemble generated samples than the AF2 predictions, and lower in most cases than the docks to the MD ensembles (Figure 5).

**Figure 5.**
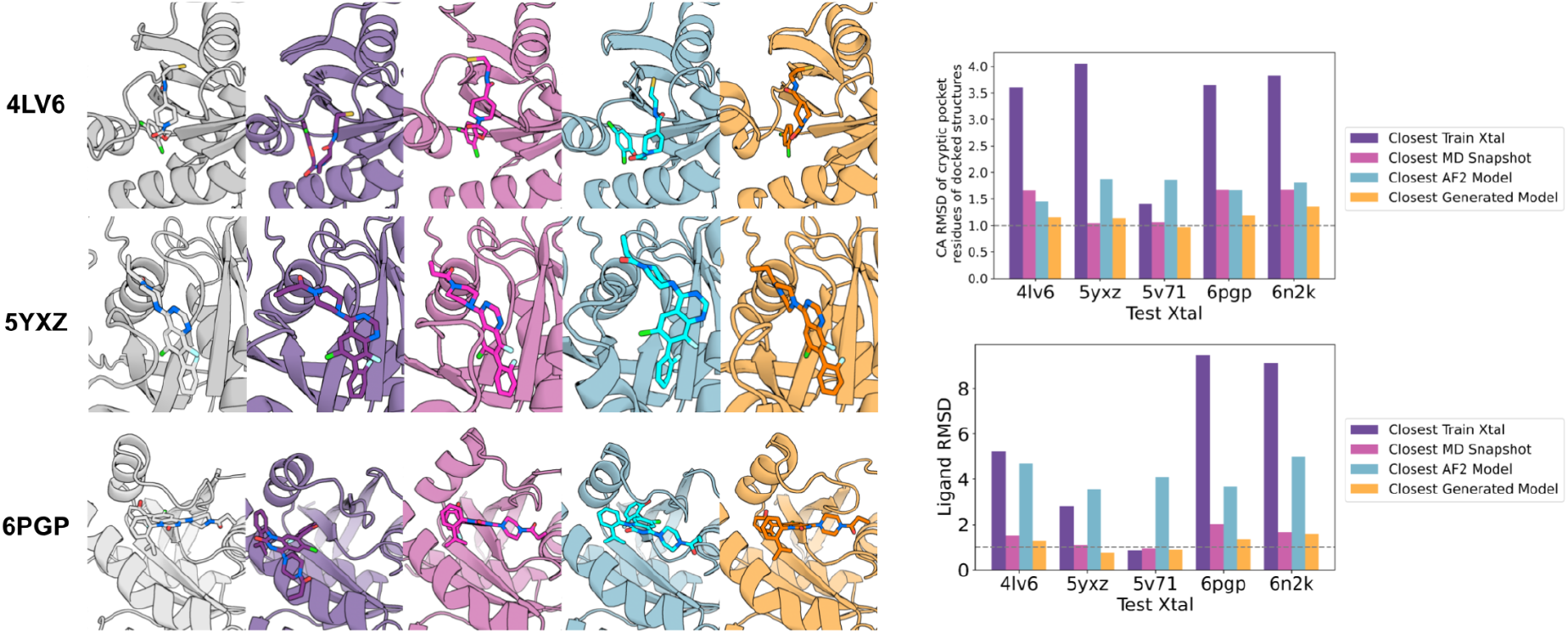
Small molecule docking into VAE generated ensembles. Ligands from held out crystal structures were docked into protein conformers using GA-ligand dock. Left: the held out crystal structure complex (column 1) and the closest docked complex (in terms of RMSD over the ligand) among the training set crystal structures (column 2), the MD snapshots (column 3), the AlphaFold models (column 4), and the VAE ensembles (column 5). The closest RMSDs of C-alpha coordinate RMSD of the cryptic pocket of docked structures and lowest RMSD over ligand atoms (ligand RMSD) are indicated on the bar charts on the right.

## Discussion

Our VAE based sampling approach enables extrapolation from combinations of MD simulation snapshots starting from multiple known crystal structures to generate ensembles of conformers more closely sampling held out crystal structures. The ensembles sample alternative ligand binding site geometries sufficiently accurately to enable docking of small molecule ligands. Our approach provides a way to generalize from multiple classical MD simulation trajectories from different crystal structure starting points to generate an effectively unlimited number of plausible samples with very low computational cost. We go beyond previous studies using VAEs to model the space sampled by MD simulations by taking advantage of the sophisticated understanding of protein sequence-structure relationships implicit in the AF2 and RF deep neural networks in two ways: first, we use the AF predicted distance distributions to focus the latent space sampling on regions consistent with the amino acid sequence, and second, we use RF to generate 3D coordinates from the output distance maps which ensures physical realism and local sequence-structure compatibility.

There are clear paths forward for improving our approach. First, the reconstruction error of ∼1Å for the known crystal structures is reasonable, but the challenge is that the differences between many of the different conformations are also of this order, limiting the ability of our approach to really precisely sample alternative states. VAE architectures with still lower reconstruction errors would improve our method, as could fine-tuning the trained VAE on the FAPE loss (we did not observe this in preliminary tests, but this warrants further exploration). Second, while the AF2 CCE metric provides a reasonable guidepost, AF2 is trained to generate single structures, and hence the use of this measure to guide sampling could limit diversity. Better results could be obtained by minimizing towards a predicted ensemble of structures for a given target or subsampling the target MSA for RoseTTAFold structure generation (Meller et al., 2023) to introduce more diversity in output structures. Despite these limitations, our results show the utility of deep generative models for modeling the conformational ensembles that determine protein function and drugability.

## Methods

### 1. Input data setup and incremental learning

For the input dataset, we began by selecting distinct K-Ras conformations deposited in the PDB that are at least an angstrom away from each other as our ‘training set crystal structures’. In addition to the RMSD cut-off filter, we also selected conformations that had a deposited / known inhibitor. We selected 20 K-Ras structures with these criteria. We ran MD simulations for 10 ns starting with each K-Ras crystal structure and selected every 50 ps snapshots from 5 independent trajectories, giving a total of 1000 MD snapshots for each starting structure. AMBER19SB force field (Tian et al., 2020) with TIP3P water model (Jorgensen et al., 1983) was used in a periodic boundary box. Langevin dynamics was run at a constant temperature of 300K and pressure of 1 atm. For each target crystal, the training data consisted of MD snapshots of the training set crystal structures that were at least an angstrom away from it. The final 20 K-Ras conformations that we chose were: 4DSO, 5XCO, 5YXZ, 6PGP, 7EWB, 8AFD, 8DNI, 4LV6, 4L9W, 5V9O, 6B0V, 6N2K, 6P8W, 7RT1, 7U8H, 4Q21, 5V71, 5E95, 6H46 and 7C40. All 3D structures were converted to 2D template features from RoseTTAFold (Baek et al., 2021) which consists of Cb distances and orientations. We chose to use the raw distance and orientation values for training the model for a more interpretable latent space.

After the first round of training using only MD snapshots as the training data, we then generated 3000 samples from the latent space that were optimized for the score metric and passed the diversity filter (following the protocol laid out in the sampling methods section). These 3000 generated structures were then concatenated on the initial MD snapshot training set to form an ‘incremental learning’ training set of structures for the model. Using this new set, for each target, the training runs were set up again from scratch. Incremental learning in this case benefits the VAE by providing a larger and more diverse set of structures for exploration, improving representation of structural diversity, refining metric optimization, and ultimately increasing the accuracy of the generated samples to the target crystal.

### 2. Soft Introspective VAE objective and training

We found best results using a Soft-Introspective VAE architecture (Daniel et al., 2020) which has been shown to have higher output resolution than the vanilla VAE (Kingma et al., 2014). The objective function of this model, along with the traditional VAE objective function of reconstruction loss and KL divergence, has adversarial losses incorporated, like GANs (Goodfellow et al., 2014) but is trained introspectively. In the case of SI-VAEs, the encoder is the implicit ‘discriminator’ where it is induced to distinguish, through the ELBO (evidence lower bound) (Kingma et al., 2014) values that it assigns to the real and generated samples. The decoder is the ‘generator’ where it is induced to generate samples to ‘fool’ the encoder (discriminator). However, unlike GANs, the SI-VAE model does not converge to the data distribution, but to an entropy-regularized version of it (Daniel et al., 2020).

Using default parameters from Daniel *et al*., 2020, encoder was trained with the following objective (Equation 1):

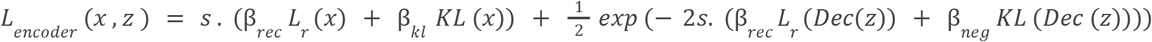

where *L*_*r*_(*x*) = reconstruction loss, *s* = 2, β _*rec*_=10,β _*kl*_=1e-3,β _*neg*_=latent dimension = 256 and Dec = trained decoder of soft-introspective VAE.

The decoder was optimized using the following objective (Equation 2):

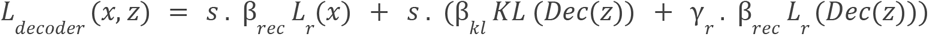

where *L*_*r*_(*x*) = reconstruction loss, *s =* 2, β _*rec*_=10,β _*kl*_ = 1e-3 and γ_*r*_ = 1.0

The reconstruction loss was the mean-squared-error loss over all distances and orientations on the decoded template features from the model. The model was optimized using individual optimizers for the encoder and decoder, both of which were initialized with Adam (β1 = 0.9, β2 = 0.999) with learning rate 1e−3, with an effective batch size of 64. The encoder and decoder were made up of 3 ResNet blocks with 2D convolutional layers and the latent space was kept at a constant of 256 dimensions for all targets.

### 3. Sampling in latent space through gradient optimization of score metric (CCE)

To obtain the optimized structures using the trained decoder, we used gradient optimization in the latent space. We first randomly sample *n* numbers from the standard Gaussian distribution (mean=0, std=1) with dimension equal to that of the latent space. The initialized latent space coordinates are set to be trainable. Each sample is then decoded into its respective template features and Cb distances are discretized through radial basis function to ensure back propagation. The score metric we chose to optimize is the minimum categorical cross-entropy (CCE) among all 5 AF2 predicted Cb distograms of the target structure and the generated Cb distances. The Adam optimizer modifies the latent space sample to minimize this score metric. This process is repeated until convergence. To ensure that diversity is maintained, the latent space coordinates are restricted to explore only *d* (=10) euclidean distance in the latent space from their initial starting coordinates. The overall goal of this exploration technique is to search the latent space to find a better solution near the initial randomly generated coordinates. The final, converged latent space coordinates are decoded into their respective template features and passed into RoseTTAFold, along with the target MSA for structural modeling.

### 4. Docking Protocol

For each docking case, the inhibitor ligand was docked to the receptor model using the protein-ligand docking method Rosetta GALigandDock (Park et al., 2021). The ligand atomic coordinates found in complex crystal structures were extracted and used to prepare for ligand docking. The ligands were protonated and the AM1-BCC partial charges were calculated using the tools provided by openbabel, Antechamber in the AMBER suite, and UCSF Chimera (Pettersen et al., 2004). The ligand information was converted to the parameter format that is compatible with the Rosetta generic potential (*GenFF* (Park et al., 2021)). The initial position of the ligand to initiate docking was determined by superimposing the complex crystal structure to the sampled protein backbone. Protein-ligand docking was performed by allowing the side chains that are within 6A from the ligand to be flexible. The receptor models were optimized in advance using Rosetta FastRelax with high constraints on each backbone. We ran 20 parallel docking runs for each receptor model and ligand pair, and the combined results were analyzed, where the best scoring generated sample was compared to best scoring models of the training set, training crystals and AlphaFold models.

## Acknowledgements and Disclosure of Funding

We would like to thank Doug Tischer, Ivan Anishchanka, Sam Pellock for helpful comments and suggestions. This work was supported by Microsoft (S.M. M.B., D.B., and generous gifts of Azure computing time), Eric and Wendy Schmidt by recommendation of the Schmidt Futures (H.P.), The Washington Research Foundation, Innovation Fellows Program (G.R.L), The Defense Threat Reduction Agency (G.R.L), The Open Philanthropy Project Improving Protein Design Fund (G.R.L), The Audacious Project at the Institute for Protein Design (D.B.), a gift from Amgen (D.B.) and the Howard Hughes Medical Institute (D.B.).

## Supplementary information

**Figure S1.**
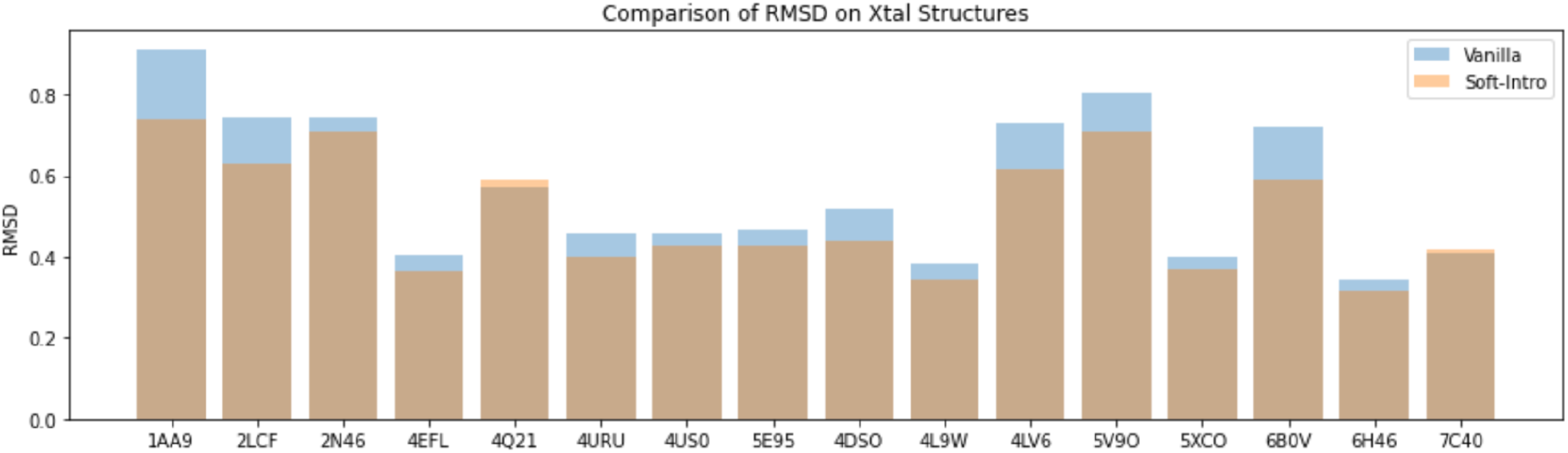
Comparison of reconstruction performance between vanilla VAE and soft-introspective VAE. Distance RMSD comparison of reconstruction of a different set of K-Ras training crystals from similarly trained vanilla VAE and soft-introspective VAE.

**Figure S2.**
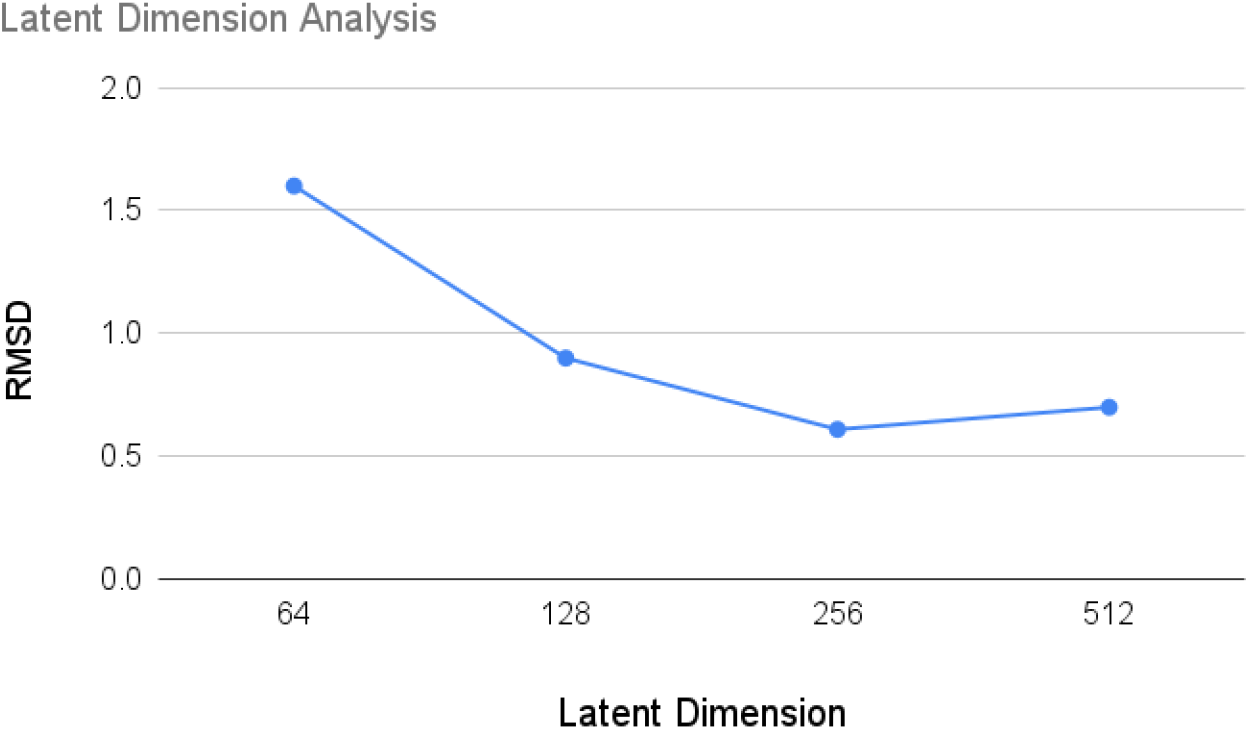
Graph illustrating the relationship between latent dimension and RMSD. Increasing the dimension (x-axis) initially leads to a significant decrease in mean RMSD calculated over training data (y-axis), indicating improved data representation. However, the graph reaches an elbow point (256 dimensions) where further dimension expansion yields diminishing returns, plateauing the RMSD reduction.

**Figure S3.**
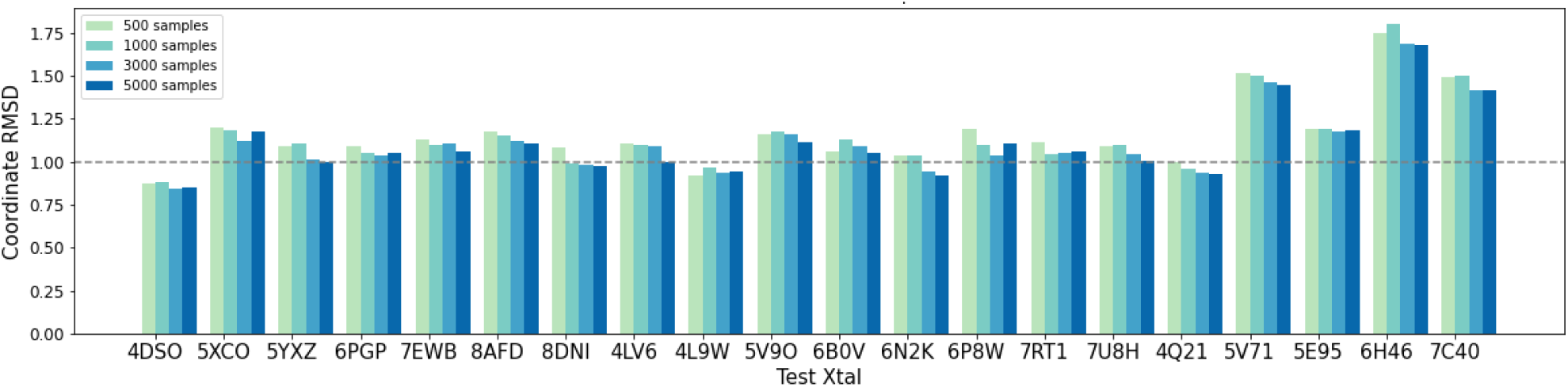
Relationship between increasing number of samples generated in latent space and closest coordinate RMSD to target. Each target is associated with a different number of samples generated in the latent space, and the corresponding Closest Coordinate RMSD to the target crystal is plotted. More samples result in lower RMSD until a threshold is reached, indicating improved accuracy.

